# A safe insect-based Chikungunya fever vaccine affords rapid and durable protection in cynomolgus macaques

**DOI:** 10.1101/2024.05.21.595029

**Authors:** Awadalkareem Adam, Courtney Woolsey, Hannah Lu, Kenneth Plante, Shannon M. Wallace, Leslie Rodriguez, Divya P. Shinde, Yingjun Cui, Alexander W.E. Franz, Saravanan Thangamani, Jason E. Comer, Scott C. Weaver, Tian Wang

## Abstract

Chikungunya virus (CHIKV), which induces chikungunya fever and chronic arthralgia, is an emerging public health concern. Safe and efficient vaccination strategies are needed to prevent or mitigate virus-associated acute and chronic morbidities for preparation of future outbreaks. Eilat (EILV)/CHIKV, a chimeric alphavirus which contains the structural proteins of CHIKV and the non-structural proteins of EILV, does not replicate in vertebrate cells. The chimeric virus was previously reported to induce protective adaptive immunity in mice. Here, we assessed the capacity of the virus to induce quick and durable protection in cynomolgus macaques. EILV/CHIKV protected macaques from wild-type (WT) CHIKV infection one year after a single dose vaccination. Transcriptome and *in vitro* functional analyses reveal that the chimeric virus triggered toll-like receptor signaling and T cell, memory B cell and antibody responses in a dose-dependent manner. Notably, EILV/CHIKV preferentially induced more durable, robust, and broader repertoire of CHIKV-specific T cell responses, compared to a live attenuated CHIKV 181/25 vaccine strain. The insect-based chimeric virus did not cause skin hypersensitivity reactions in guinea pigs sensitized to mosquito bites. Furthermore, EILV/CHIKV induced strong neutralization antibodies and protected cynomolgus macaques from WT CHIKV infection within six days post vaccination. Transcriptome analysis also suggest that the chimeric virus induction of multiple innate immune pathways, including Toll-like receptor signaling, type I IFN and IL-12 signaling, antigen presenting cell activation, and NK receptor signaling. Our findings suggest that EILV/CHIKV is a safe, highly efficacious vaccine, and provides both rapid and long-lasting protection in cynomolgus macaques.

## Introduction

Chikungunya virus (CHIKV), a reemerging mosquito-borne positive-sense RNA virus, belongs to the genus Alphavirus of the family *Togaviridae*. The virus was initially isolated in 1952 in Tanzania, and has caused epidemics in Africa, Asia, Europe, and the Americas with millions of human cases. The virus induces acute and chronic human infections with main clinical symptoms including chikungunya fever (CHIKF) and severe joint pain (1, 2). CHIKV has a 5’ capped RNA genome of 11.8 kb in length that encodes four nonstructural proteins (nsP1–4) and six structural proteins, including the capsid protein (C), envelope (E) glycoproteins (E3, E2, and E1), 6K viroporin channel, and transframe proteins (TM) (3–6). There are four major lineages of CHIKV: Asian Urban (AUL), Indian Ocean (IOL), East/ Central/South African (ESCA), and West African (WA) enzootic lineages (7). An effective vaccine is needed to prevent or reduce CHIKV-associated chronic morbidity. Multiple platforms have been used for CHIKV vaccine development, including formalin-inactivated, virus-like particles (VLP), recombinant subunit, DNA, virus-vector-based, live-attenuated, and chimeric alphavirus-based vaccines(8–17). In November 2023, Ixchiq, a live attenuated CHIKV vaccines which was developed as VLA1553 (10) was approved by the U.S. Food and Drug Administration. Despite this achievement, additional CHIKV vaccination strategies are needed.

We previously reported the generation of Eilat (EILV)-CHIKV, a chimeric virus containing the structural proteins of CHIKV (E1, 6K, E2, E3 and capsid) and the non-structural (NS) proteins (nsP1, nsP2, nsP3 and nsP4) of EILV, an insect-specific alphavirus that is defective for replication in vertebrate cells (18). Like EILV, the chimeric virus replicates well in mosquito cells, but not in vertebrate cells (19, 20). A single dose of EILV/CHIKV induces cellular and humoral immunity and protects A129 IFNAR^−/−^ mice from CHIKV-induced disease up to 9.5 months post vaccination. Furthermore, EILV/CHIKV completely protects cynomolgus macaques from CHIKF one month post vaccination (20, 21). However, the underlying mechanism and immune activation of EILV/CHIKV are not well understood. In this study, we characterized innate immune signaling and the longevity of EILV/CHIKV-induced T cells, memory B cells and antibody responses and understand immune factors correlated with rapid and durable protection following a single dose vaccination in cynomolgus macaques.

## Results

### A single dose of EILV/CHIKV protects cynomolgus macaques from wild-type (WT) CHIKV infection one year after vaccination

To determine the durability of EILV/CHIKV-induced protective efficacy, 2.5- to 3.5-year-old female cynomolgus macaques were vaccinated with a low (LD) and high dose (HD) of EILV/CHIKV. CHIKV 181/25 or PBS (mock)-vaccinated macaques were used as controls (**Fig. 1A**). Animals were bled at various time points post-vaccination (PV) to assess immunogenicity. At day 370 PV, macaques were challenged with the wild-type (WT) CHIKV La Réunion strain and monitored daily for clinical signs. None of the macaques displayed obvious signs nor swelling at the injection site. While mock-vaccinated macaques showed fever which peaked around day 1 post challenge (PC); CHIKV 181/25 and EILV/CHIKV-vaccinated macaques displayed normal temperature (**Fig. 1B**). Viremia peaked at day 1 PC and was 3 log_10_ higher in the mock group than EILV/CHIKV (LD and HD) and CHIKV 181/25-vaccinated macaques. At day 4 PC, viremia was no longer detectable by plaque assay in the vaccinated groups but remained high in the mock group. There were significantly lower levels of viral RNA in the EILV/CHIKV HD group than the mock (**Fig. 1C, and Supplementary Fig. 1A-B**). When compared to each other, histopathological examination revealed that the mock, CHIKV 181/25, and one of the EILV/CHIKV LD-vaccinated macaques had equal incidence and severity of decreased lymphocytes in the lymphoid nodules of spleen tissues at day 14 PC, which was not observed in the macaques-vaccinated with a HD of EILV/CHIKV (**Supplementary Fig. 1C).** Overall, these results suggest that EILV/CHIKV protects macaques from CHIKV-induced fever (CHIKF) and viremia one year PV.

**Figure 1.**
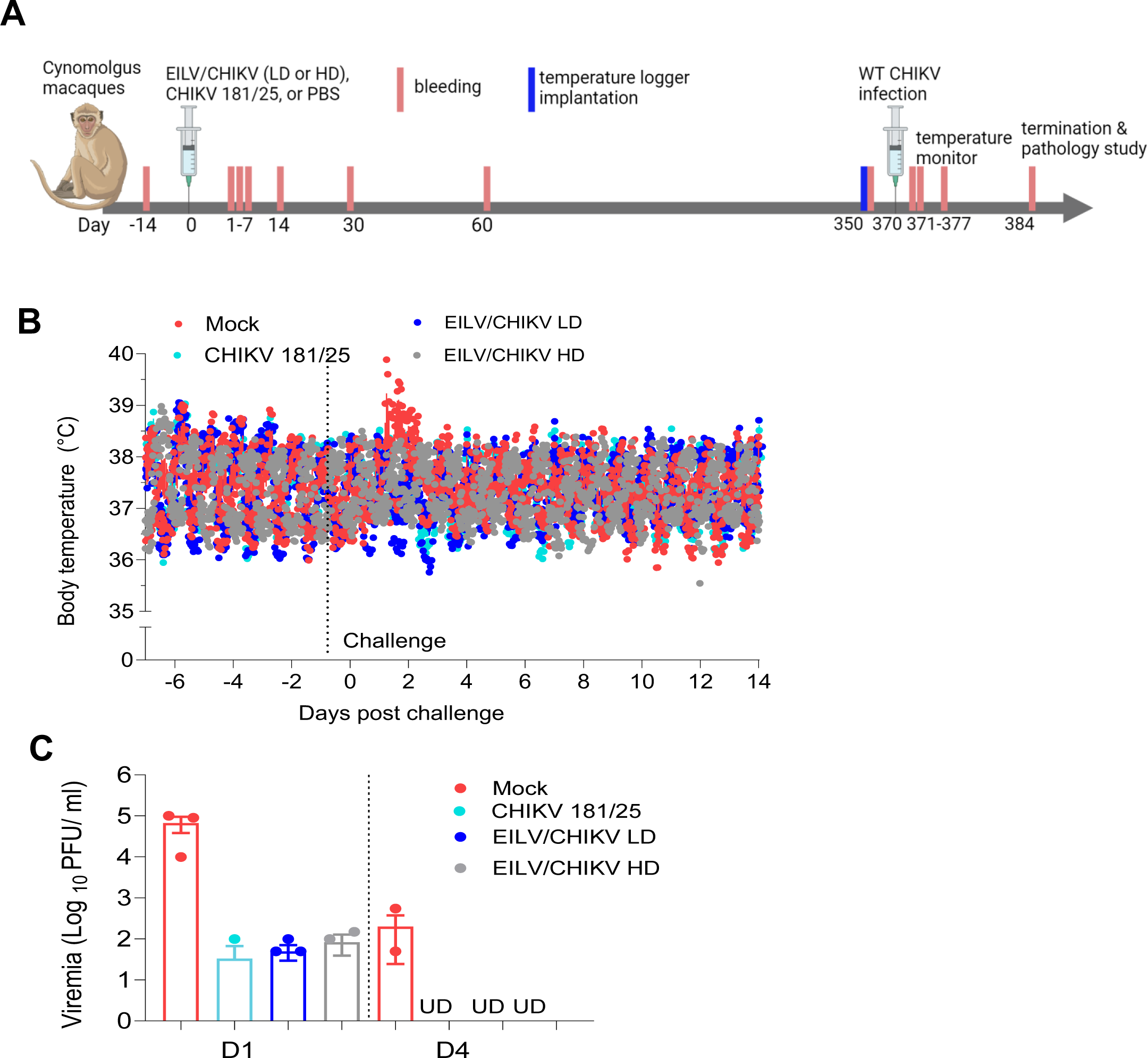
EILV/CHIKV protects cynomolgus macaques from WT CHIKV infection one year after a single dose vaccination. NHPs were vaccinated with a low (LD) or high doses (HD) of EILV/CHIKV, CHIKV 181/25 or PBS (mock). At day 350, all animals were implanted with electronic data loggers as temperature sensor and challenged subcutaneously with 1.0 × 10^5^ PFU of WT La Reunion strain of CHIKV at day 370. **A.** Study design and vaccination timeline. **B.** Body temperature changes were recorded every 15 min and reported as mean ± standard error of the mean (SEM) starting 7 days before to 14 days after infection with WT CHIKV strain La Réunion. **C**. Viremia were measured by plaque assays at indicated days post-challenge (PC). UD: Undetectable. Data are presented as means ± SEM.

### EILV/CHIKV triggers potent adaptive immunity in a dose-dependent manner and preferentially induces durable T cell immunity

To understand the underlying mechanisms of immune protection, we performed transcriptome analysis of PBMCs of EILV/CHIKV (HD), CHIKV 181/25 and mock groups at day 4 PC. Principal component analysis (PCA) indicated dimensional separation of the mock and vaccinated samples (**Fig. 2A**). EILV/CHIKV and CHIKV 181/25 samples clustered together, indicating similar overall expression profiles. To identify protective signatures specific for each vaccine, we focused on differences between EILV/CHIKV and CHIKV 181/25 vaccination for our differential expression analysis (**Fig. 2B**). EILV/CHIKV versus CHIKV 181/25 subjects expressed higher levels of T cell-associated transcripts (*CCR7*, *CD28*, *CD5*, *CD3G*, *CD8A*). EILV/CHIKV vaccination also robustly activated pathways involved in T and B cell signaling including “co-stimulation by the CD28 family”, “CD40 signaling”, “April mediated signaling”, “B cell activating factor signaling”, and “T helper (h)1” pathways. ). Molecules (*EIF2AK2*, *IKBKB*, *MAP2K4*, *MAP3K1*, *MAPK1*, *TNFAIP3*, *TRAF1*) involved in the “Toll-like Receptor (TLR) Signaling” pathway were also upregulated in subjects immunized with EILV/CHIKV (**Fig. 2C)**. Compared to the CHIKV 181/25 group, macaques immunized with EILV/CHIKV also expressed higher levels of transcripts involved in immunoregulation (e.g., IL-10, CTLA-4, PD-1) (**Fig. 2D**). To track changes in circulating cell populations, we employed nSolver-based profiling of immune cell types. For the EILV/CHIKV, we detected an expected increase in T cell, CD8^+^ T cell, CD45, and B cell transcriptional quantities (**Fig. 2E**). Conversely, except for B cells, these cell populations were decreased in the CHIKV 181/25 group. These results suggest EILV/CHIKV vaccination induces a mixed T and B cell/immunoregulatory response; whereas B cell activation was more notable following CHIKV 181/25 vaccination.

**Figure 2.**
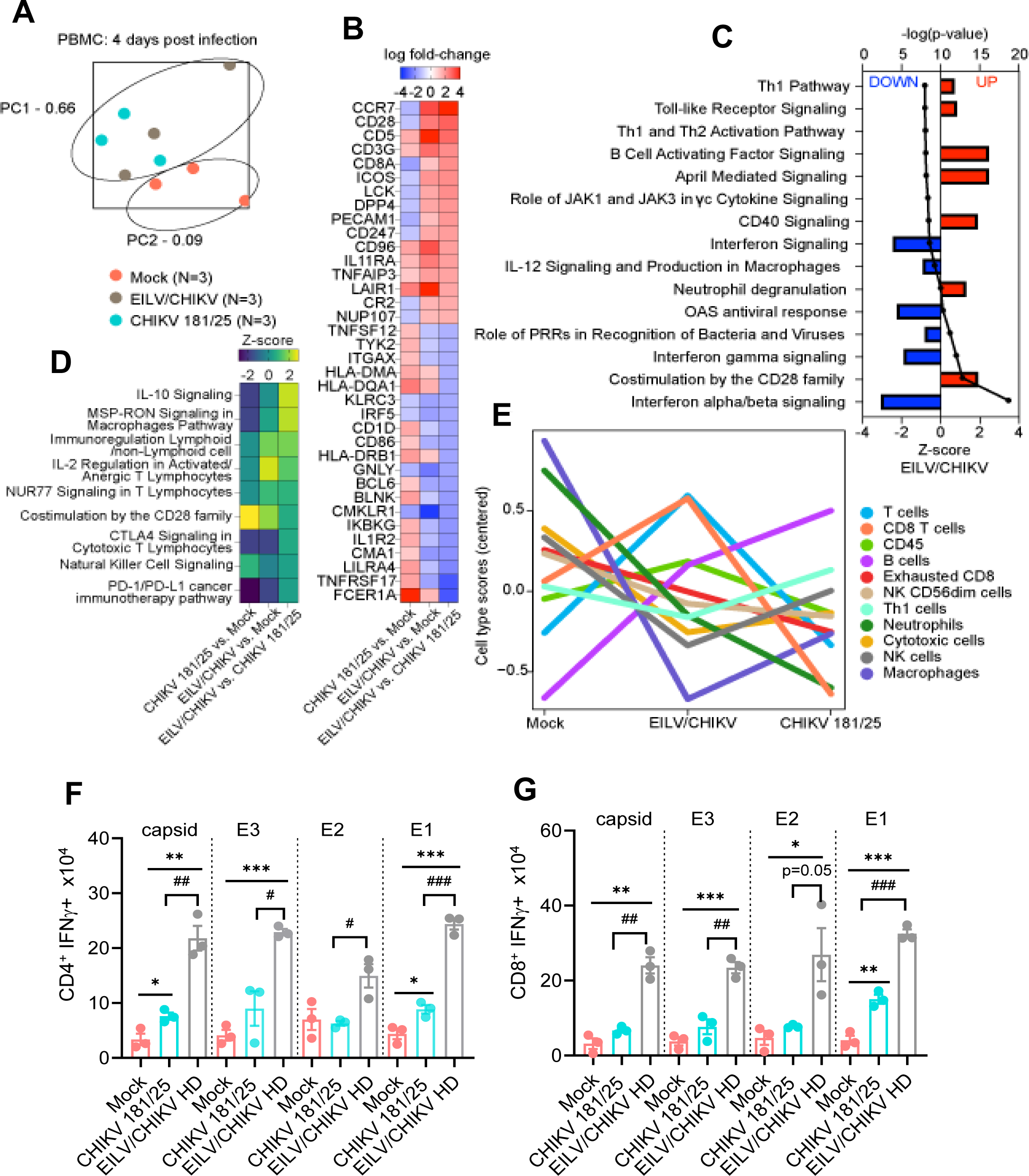
Transcriptome and T cell functional analysis of PBMCs one year after vaccination and challenge of cynomolgus macaques. **A-E.** Transcriptional changes of PBMCs of vaccinated macaques and mock controls at day 4 PC. **A.** Principal component analysis (PCA) of all normalized transcripts to visualize the relatedness of samples via dimensional reduction. Each dot represents an individual RNA sample for mock (pink; n=3), EILV/CHIKV (gray; n =3), and CHIKV 181/25 (aqua; n=3) groups. Samples that cluster are considered overall more similar, whereas distant samples are considered overall more distinct. **B.** Heatmap of the most differentially expressed mRNAs in each group. Transcripts with a Benjamini-Hochberg adjusted p-value < 0.05 were sorted by log fold change values for EILV/CHIKV-vaccinated compared to CHIKV 181/25-immunized subjects. Red indicates increased expression; blue indicates decreased expression. **C.** Bar plot of the most significant signaling pathways in EILV/CHIKV-vaccinated compared to mock-immunized subjects. Any differentially expressed transcripts with a Benjamini-Hochberg corrected *P* < 0.1 were deemed significant for enrichment. Pathways were sorted by statistical significance (a higher −log (*P*) indicates higher significance). Red indicates increased expression; blue indicates decreased expression. **D**. Heatmap of the most upregulated pathways in each group; samples were sorted by z-score for EILV/CHIKV-vaccinated compared to CHIKV 181/25-immunized subjects. Yellow/green indicates increased expression, blue/purple indicates decreased expression. **E.** Dot plots depicting predicted cell type trend scores for mock (pink; n=3), EILV/CHIKV (gray; n=3), and CHIKV 181/25 (aqua; n=3) groups. Values were derived from transcriptional signatures indicative for a particular cell subset. **F-G.** PBMCs of day 350 vaccinated NHPs were cultured *ex vivo* with CHIKV capsid, E3, E2 and E1 peptide pools for 6 h, and stained for IFN-γ, CD3, CD4, or CD8. Total number of IFN-γ^+^ CD4^+^ and CD8^+^ T cells is shown. \*\*\**P* < 0.001, \*\**P* < 0.01, or \**P* < 0.05 compared to mock. ^####^*P* < 0.001, ^##^*P* < 0.01, or ^#^*P* < 0.05 compared to CHIKV 181/25.

To verify results of the transcriptome analysis, we next performed *ex vivo* intracellular cytokine staining (ICS). PBMCs of the vaccinated macaques were treated with individual peptide pools of CHIKV capsid, E3, E2 and E1 proteins. At day 30 PV, a HD of EILV/CHIKV induced strong CD4^+^IFNγ^+^ and CD8^+^IFNγ^+^ responses reactive to all 4 peptide pools. Compared to EILV/CHIKV HD, CHIKV 181/25 induced lower magnitude of CD4^+^IFNγ^+^ and CD8^+^IFNγ^+^ responses reactive to capsid, E3, and E1 peptide pools (**Supplementary Fig. 2A**). At day 350 PV, CD4^+^IFNγ^+^ and CD8^+^IFNγ^+^ responses reactive to all structural proteins remained strong in the EILV/CHIKV HD group, but CHIKV 181/25 showed diminished levels of capsid and E1-specific CD4^+^IFNγ^+^ and CD8^+^IFNγ^+^ responses compared to the EILV/CHIKV HD group (**Fig. 2F-G**). Notably, the EILV/CHIKV LD group had comparable levels of T helper (h)-1 response compared to the CHIKV 181/25 group at day 7 PV but much lower Th1-responses against the four structural protein peptide pools at day 30 PV than CHIKV 181/25 (**Supplementary Fig. 2B).)**. These findings indicate dose-dependent responses in EILV/CHIKV-vaccinated subjects.

Next, we utilized conventional B-cell ELISpot to measure CHIKV-specific memory B cell (MBC)s in PBMCs of vaccinated macaques and mock controls. Because circulating MBCs do not actively secrete antibodies, we stimulated the PBMCs with the TLR7/8 agonist, R848, and rIL-2 *in vitro* for 5 days to convert MBCs into antibody secreting cells (ASC)s. Ig capture antibody, CHIKV structural protein peptide pools, CHIKV VLP, or EILV/CHIKV were all used as antigens to detect total ASCs, and CHIKV-specific MBCs (**Fig. 3A, Supplementary Fig. 3A)**. At day 30 PV, EILV/CHIKV HD and CHIKV 181/25 both induced higher frequencies of CHIKV-specific MBCs than the mock group. EILV/CHIKV HD induced similar levels of CHIKV-specific MBCs compared to the CHIKV 181/25 group, whereas EILV/CHIKV LD group had a lower frequency of IgG^+^-expressing MBCs. No differences were noted between the two groups at day 7 PV (**Supplementary Fig. 3B).** At day 350 PV, the CHIKV 181/25 group showed significantly higher frequency of IgG^+^ expressing MBCs than the EILV/CHIKV HD group (**Fig. 3B-C**). All vaccinated groups showed higher levels of neutralization antibodies at day 350 PV and at day 4 PC than the mock group (**Fig. 3D**). EILV/CHIKV LD induced comparable levels of neutralization titers as CHIKV 181/25 at days 14, 30, and 384 except at day 60, which appears to be lower compared to the latter (**Supplementary Fig. 3C**). The magnitude of E2-binding IgG antibody responses were significantly reduced in EILV/CHIKV (LD and HD) groups compared to those of the CHIKV 181/25 group (**Fig. 3E**). To determine the protective effects of EILV/CHIKV-induced humoral immunity, we performed adoptive transfer into AB6 mice of pooled sera from CHIKV-181/25 or EILV/CHIKV HD groups at day 350 PV followed by infection with a lethal dose of WT CHIKV. There were increased survival rates and reduced weight loss in the mice transferred with sera of both vaccinated NHP groups. Mice transferred with immune sera of CHIKV-181/25-vaccinated NHPs had higher survival (50 % vs. 20%) compared to mice transferred with EILV/CHIKV HD sera, suggesting protection is partially mediated by humoral responses, particularly for CHIKV-181/25 (**Fig. 3F-G**). Overall, EILV/CHIKV induced T cell and humoral immune responses in a dose-dependent manner. Compared to CHIKV 181/25, EILV/CHIKV HD preferentially induced potent and durable T cell immunity against all four structural proteins.

**Figure 3.**
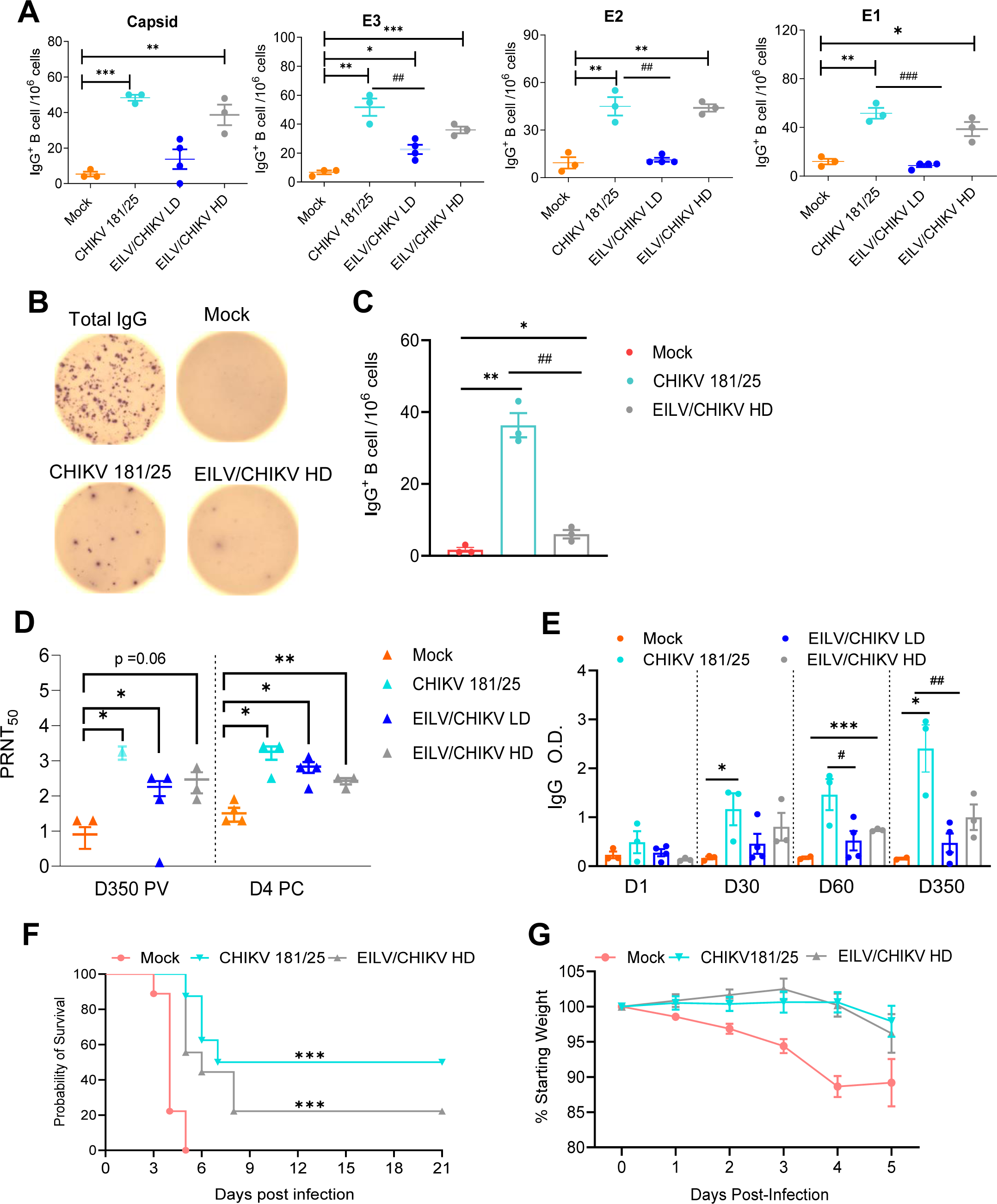
Memory B cell (MBC) and antibody responses in EILV/CHIKV-vaccinated macaques. **A-C.** CHIKV-specific MBC responses by ELISPOT analysis. PBMCs of day 30 (**A**) and day 350 (**B-C**) vaccinated NHPs were stimulated for 7 d with R848 plus rIL-2 and seeded onto ELISPOT plates coated with CHIKV capsid, E3, E2 and E1 peptide pools (**A**), total IgG or CHIKV recombinant E2 protein (**B-C**). **A & C**. Frequencies of CHIKV-specific antibody secreting cells (ASC)s per 10^6^ input cells in MBC cultures from the subject. **B.** Images of total IgG-ASCs, CHIKV peptide-specific or CHIKV E2 protein-specific MBCs, are shown. **D.** Serum neutralizing activity against CHIKV 181/25 was measured by a plaque reduction neutralization test (PRNT). **E.** CHIKV E2 binding IgG responses at indicate time points by ELISA. **F-G.** Passive immunization study. Pooled sera collected at day 350 PV of macaques were diluted 1:2 in PBS and transferred to 6-week-old AB6 mice 24 h before infection with a LD100 dose of WT CHIKV. Mice were monitored daily for morbidity. **F**. Survival rate. **G**. Percent weight loss compared to prior infection. \*\*\**P* < 0.001, \*\**P* < 0.01, or \**P* < 0.05 compared to mock. ^####^*P* < 0.001, ^##^*P* < 0.01, or ^#^*P* < 0.05 compared to CHIKV 181/25.

### EILV/CHIKV is a safe vaccine and provides rapid protection against WT CHIKV challenge

To determine whether purified EILV/CHIKV, which was produced in mosquito cells, can trigger hypersensitive reactions upon vaccination of animals sensitized to mosquito bites, we next performed a guinea pig sensitization study. Guinea pigs were repeatedly exposed to fifty *Ades albopictus* mosquito bites every two weeks for four times, then at day 43 were vaccinated with purified EILV/CHIKV at the mosquito-exposed sites. *Aedes albopictus* salivary gland extract (SGE)- and PBS (mock)-injected guinea pigs were used as positive and negative controls (**Supplementary Fig. 4A**). SGE-injected guinea pigs showed quick skin reactions with visible swelling 30 min to 2 h after inoculation, in comparison to the mock-injected animals; EILV/CHIKV-inoculated animals showed no obvious skin reactions. (**Supplementary Fig. 4B-C).** Next, to determine whether EILV/CHIKV provides rapid protection against WT CHIKV infection. Eight 2.5- to 3.5-year-old male cynomolgus macaques were immunized i.m. with EILV/CHIKV. As the HD EILV/CHIKV proved to have stronger immunogenicity than the LD EILV/CHIKV in the long-term study, here, we used a HD of EILV/CHIKV for vaccination. The inactivated WT CHIKV- or PBS (mock)-vaccinated macaques were used as controls. At day 6 PV, all animals were challenged with WT CHIKV strain La Réunion (**Fig. 4A**). Fever peaked around day 2 for the CHIKV-infected mock group and remained high even at day 3 PC (**Fig. 4B**). Macaques-vaccinated with the inactivated WT CHIKV or EILV/CHIKV exhibited normal and baseline body temperatures throughout the study. Viremia titers more than 3 log_10_-fold lower in the inactivated WT CHIKV and EILV/CHIKV-vaccinated animals compared to the mock at days 1 and 2 PC. At day 4 PC, viral titers were diminished 3 log_10_-fold lower in the mock group but were no longer detectable in the two vaccinated groups.. Viral RNA levels were significantly reduced in the EILV/CHIKV group compared to the mock (**Fig. 4C-D**). Thus, EILV/CHIKV appears safe and triggers rapid protection against WT CHIKV infection.

**Figure 4.**
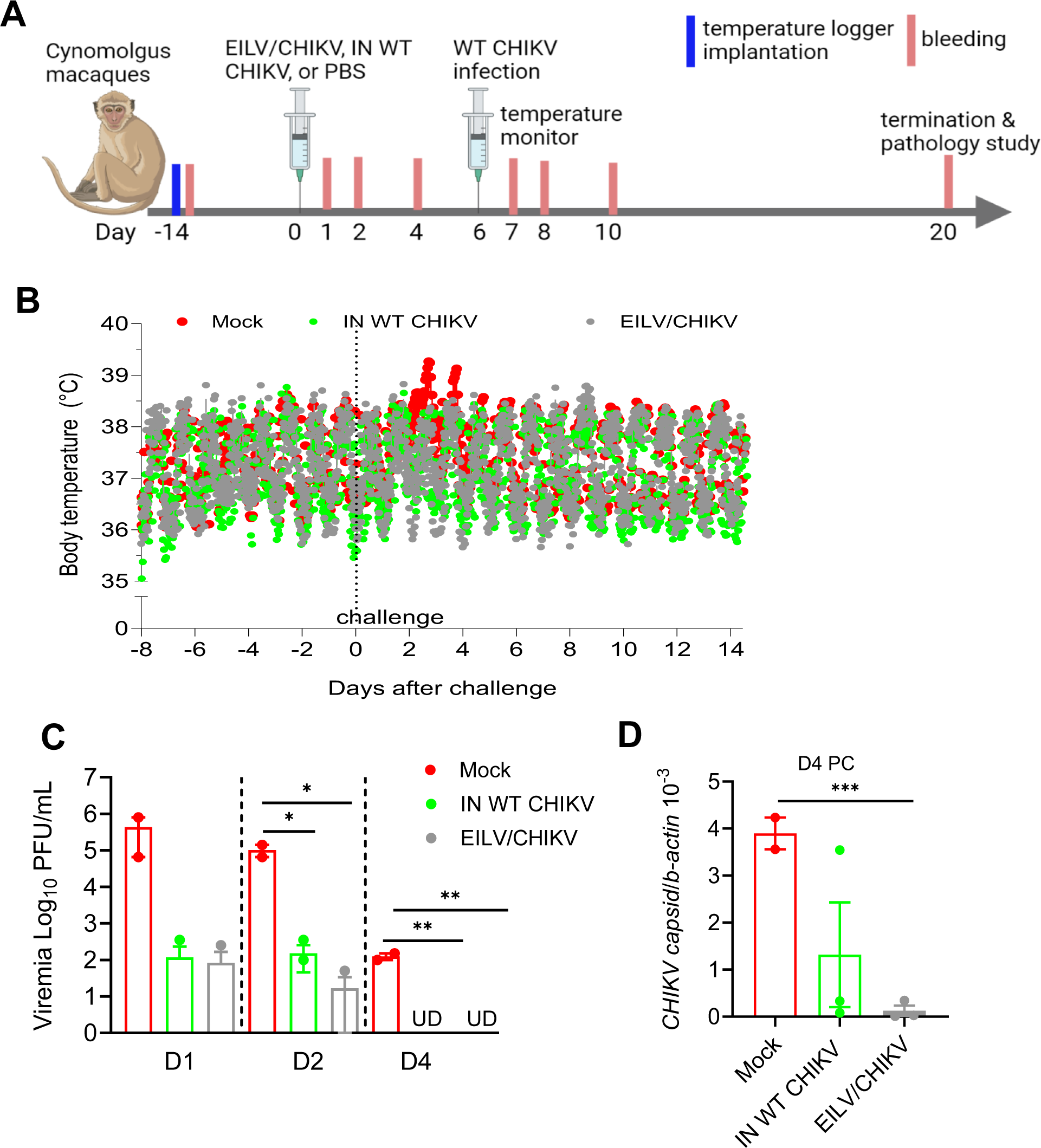
Rapid protection by EILV/CHIKV vaccination of cynomolgus macaques. NHPs were implanted with electronic data loggers as temperature sensor and bled 14 days before vaccination. NHPs were vaccinated with 1.3×10^8^ PFU EILV/CHIKV (n =3), 5×10^5^ PFU inactivated WT CHIKV (n=3) or PBS (mock, n=2). At day 6 PV, NHPs were challenged subcutaneously with 10^5^ PFU of WT La Reunion strain of CHIKV. **A.** Study design and vaccination timeline. **B.** Body temperature changes were recorded every 15 min and reported as hourly mean ± SEM starting 8 days before to 14 days after the challenge with WT CHIKV. **C-D**. Viremia were measured by plaque assays (**C**) or Q-PCR (**D**) at indicated days PC. Undetectable: UD. ** *P* < 0.01 compared to mock group. Unpaired, 2-tailed Student’s t test was used to determine the differences. Data are presented as means ± SEM.

### EILV/CHIKV stimulates a rapid type I interferon (IFN) response, robust natural killer (NK) cell expansion, and potent adaptive immunity within a week post-vaccination

To understand the immune mechanisms of rapid host protection, we analyzed transcriptomes of PBMCs of EILV/CHIKV- or mock-immunized macaque at day 7 PV. PCA revealed distinct transcriptional profiles for mock) and EILV/CHIKV groups (**Fig. 5A**). During the vaccination phase, subjects administered EILV/CHIKV expressed higher levels of transcripts associated with NK and T cell differentiation and cytotoxicity (*KLRK1*, *GZMB*, *EOMES*, *GZMH*, *GNLY*) (**Fig. 5B**). *HLA-DPA1* was also highly expressed in EILV/CHIKV subjects and encodes for an MHC class II protein that plays a central role in adaptive immunity by presenting peptides derived from extracellular proteins. Eomes is predominantly expressed in immature NK cells and regulates the differentiation of CD8^+^ T cells into effector and memory cells; this molecule also regulates the thymic differentiation of NK T cells and their commitment to a memory KLRG1^+^iNKT phenotype in the periphery. We also observed a general increase in expression of interferon (IFN)-related and Th1-associated transcripts, although these narrowly failed to reach statistical significance: log FC 1.1 *IFNG* (*P* =0.1898); 1.19 *IFNGR1* (*P* =0.4269); 1.54 *IFNGR2* (*P* =0.1339); 1.19 *IL12RB1* (*P* =0.1883); 1.34 *IL12RB2* (*P* =0.1854); 1.43 *JAK1* (*P* =0.1699); 1.38 *JAK2* (*P* =0.2142); 1.36 *STAT1* (*P* =0.0856); and 1.51 *STAT4* (*P* =0.2214) (**Table S1**). However, pathway analysis demonstrated an increase in IFN and IL-12 signaling (**Fig. 5C**). The most significantly upregulated pathways in vaccinated subjects included NK cell signaling, Th1, and Th2 pathways. B cell receptor signaling pathways were also upregulated (e.g., “signaling by the BCR”, “April mediated signaling, B cell activating factor pathway”, **Fig. 5D**). Consistent with our differential expression and pathway enrichment analyses, digital cell quantitation indicated higher predicted quantities of NK, Th1, cytotoxic T cells, and B cells. Lower predicted quantities of neutrophils were noted in vaccinated versus mock subjects.

**Figure 5.**
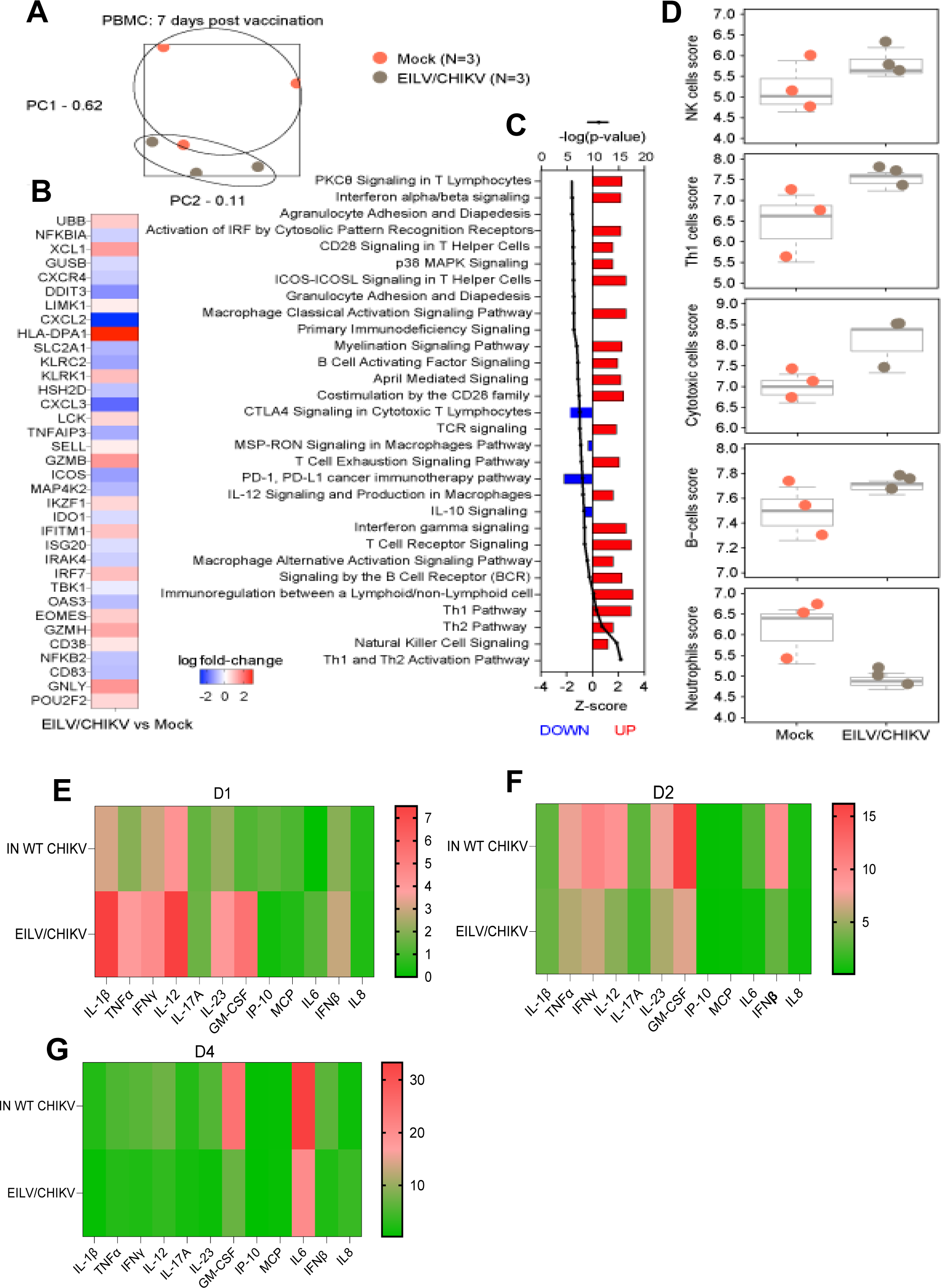
Transcriptional changes and sera cytokine production in vaccinated macaques. **A-E**. Transcriptional changes of PBMCs in EILV/CHIKV-vaccinated macaques and mock controls at day 7 PV. (**A**) PCA of all normalized transcripts to visualize the relatedness of samples via dimensional reduction. Each dot represents an individual RNA sample for mock (pink; n=3) and EILV/CHIKV (gray; n=3) groups. Samples that cluster are considered overall more similar, whereas distant samples are considered overall more distinct. (**B**) Heatmap of the most differentially expressed mRNAs in EILV/CHIKV-vaccinated compared to the mock-immunized subjects; sorted by statistical significance (Benjamini-Hochberg adjusted *P* < 0.05). Red indicates increased expression, white indicates no change in expression, blue indicates decreased expression. (**C**) Bar plot of the most significant signaling pathways in EILV/CHIKV-vaccinated compared to mock-immunized subjects. Any differentially expressed transcripts with a Benjamini-Hochberg corrected *P* < 0.1 were deemed significant for enrichment purposes. Pathways are sorted by statistical significance (a higher −log(p-value) indicates higher significance). Red indicates increased expression, blue indicates decreased expression. (**D**) Dot plots depicting predicted cell type trend scores for mock (pink; n=3) and EILV/CHIKV (gray; n=3) groups. Values were derived from transcriptional signatures indicative for a particular cell subset. **E-G.** Sera cytokines were measured by nonhuman primate inflammation 13-plex kits at indicated time points PC. Data are presented as fold increases compared to the mock group.

We next measured the production of serum cytokines in the vaccinated macaques and controls. At day 1 PC, the EILV/CHIKV group showed higher levels of IFN-β, IL-12, IL-23, GM-CSF, and proinflammatory cytokines (IL-1β, TNF-α, and IFN-γ) than the inactivated WT CHIKV group. At days 2 and 4 PC, the cytokines were higher in the inactivated WT CHIKV group than EILV/CHIKV group (**Fig. 5E-G**). Thus, EILV/CHIKV primed NHPs to induce quicker and higher type I IFN, IL-12, IL-23, GM-CSF, and proinflammatory cytokine responses than the inactivated WT CHIKV group. Interestingly, EILV/CHIKV LD vaccination also induced increased levels of expression of IL-1β, IL-6, IL-12, TNF-α and TGF-β either higher or comparable than the CHIKV 181/25 group at days 1 and 2 PV (**Supplementary Fig. 6)**. These results together with the transcriptome analysis support EILV/CHIKV as a potent inducer of innate cytokine responses. At day 4 PC, NK cell but not NK T cell expansion was noted in both the inactivated WT CHIKV and EILV/CHIKV groups and was more than 63% higher in the latter group (**Fig. 6A-B**). Furthermore, EILV/CHIKV-vaccinated macaques showed a trend of stronger neutralization titers and higher E2 binding IgG responses at day 4 PC (**Fig. 6C-D**). Lastly, both the inactivated WT CHIKV and EILV/CHIKV-vaccination boosts Th-1 responses reactive towards all structural proteins at day 4 PC compared to the mock group (**Fig. 6E**). Overall, EILV/CHIKV induces quicker type I IFN and proinflammatory cytokine responses, more robust NK cell expansion and neutralization antibody titers, and comparable levels of Th1 responses compared to the inactivated WT CHIKV in response to WT CHIKV infection one week PV.

**Figure 6.**
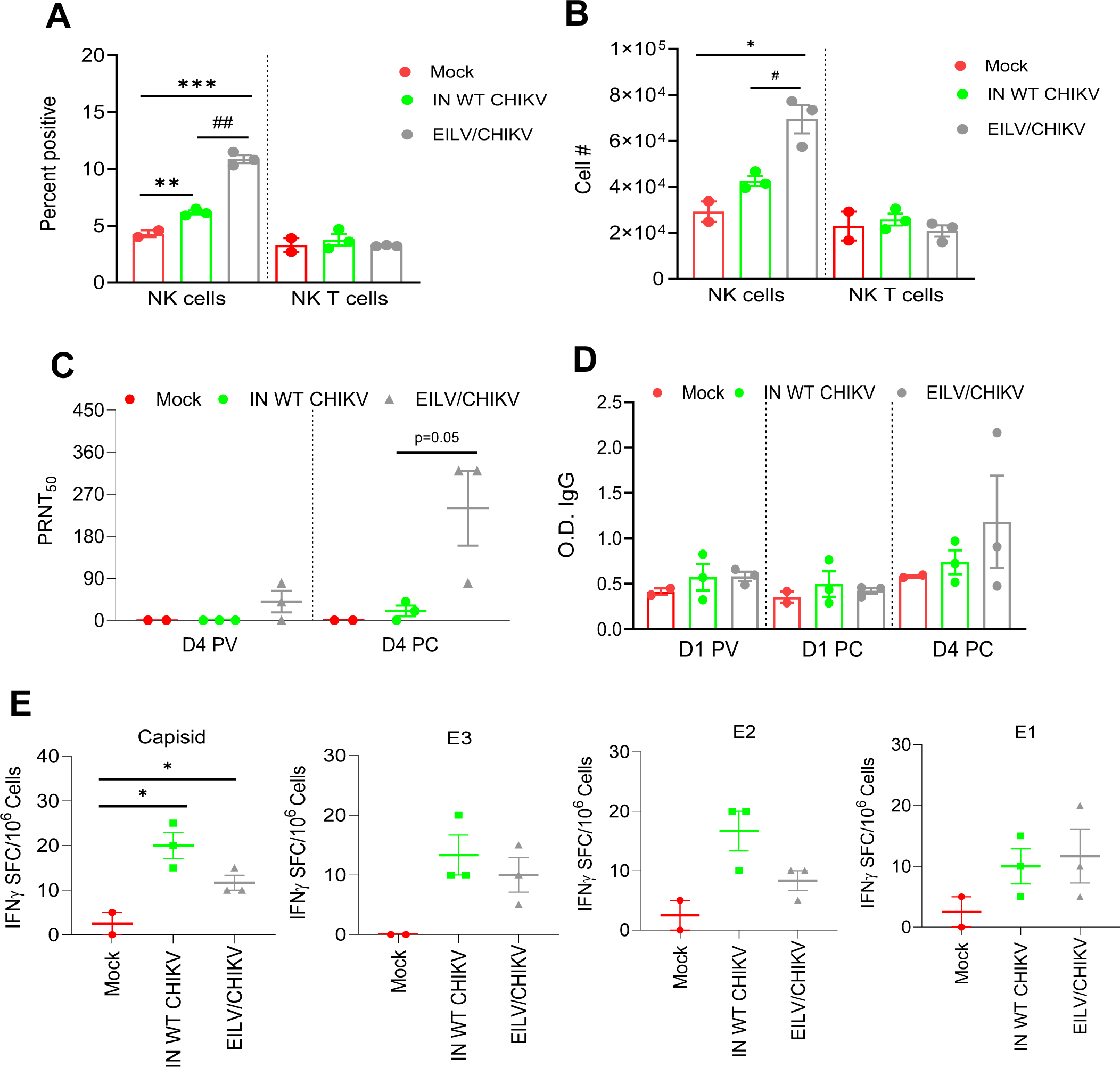
EILV/CHIKV-induced protective innate and adaptive immunity in macaques. **A-B.** NHPs were vaccinated with EILV/CHIKV, inactivated WT CHIKV or PBS (mock). At day 6 PV, NHPs were challenged subcutaneously with 10^5^ PFU of the WT La Reunion strain of CHIKV. At day 4 PC, cells expressing NK cell markers were analyzed. Percent positive (**A**) or total number (**B**) of NK or NK T cells in PBMCs are shown. **C.** Serum neutralizing activity against CHIKV 181/25 was measured by PRNT. **D.** CHIKV E2 binding IgG responses at indicate time points by ELISA. **E.** ELISPOT quantification of peripheral T cell responses. PBMCs of macaques collected at day 4 PC were stimulated with CHIKV capsid, E3, E2 and E1 peptide pools for 24 h. SFCs were measured by IFN-γ ELISPOT. Data are shown as # of SFC per 10^6^ cells. \*\**P* < 0.01, or \**P* < 0.05 compared to mock. ^##^*P* < 0.01 compared to CHIKV 181/25.

## Discussion

EILV/CHIKV induces robust adaptive immunity in mice and protects host from WT CHIKV-induced diseases 9.5 months post vaccination (20). The underlying immune mechanisms of host protection have not been clearly defined. Here, we evaluated the long-term protective efficacy of EILV/CHIKV vaccination in cynomolgus macaques and assessed its immunogenicity longitudinally. A single dose of EILV/CHIKV protected cynomolgus macaques against CHIKF one year after vaccination. Transcriptome analysis of PBMCs revealed the induction of multiple immune pathways, including TLR, T cell, and B cell signaling pathways in macaques 1 year post EILV/CHIKV vaccination followed by WT CHIKV challenge. T/B cell functional analysis supported these findings and further suggests that EILV/CHIKV-stimulates adaptive immunity in a dose-dependent manner.

Adaptive immunity including T cells, B cells, and antibody responses provide durable and specific responses against viral infection. Mature B and T cells, and B cell-mediated neutralization antibody responses contribute to host protection against CHIKV infection (22–24). Among T cells, CHIKV-specific CD8^+^ T cells are known to protect mice from WT CHIKV induced footpad swelling, to reduce inflammation, (25) and to accelerate viral clearance in lymphoid tissues (26, 27). Although EILV/CHIKV and the live attenuated CHIKV 181/25 both provided durable protection against CHIKF, the two vaccines induced differential T and B cell immune responses. EILV/CHIKV (HD) induced superior T cell immunity that was sustained 1 year after vaccination; whereas the frequency of CHIKV-specific memory B cells and E2-binding antibodies declined after 1 year, though comparable levels of Nab titers were sustained. In contrast, CHIKV 181/25, induced potent and durable memory B cell and CHIKV-specific antibodies as reported (23). However, it induced much lower T cell responses, which diminished at one year PV. Notably, passive immunization studies with immune sera of CHIKV 181/25 and EILV/CHIKV groups one year PV confirmed the differences in humoral immunity between the two groups. Furthermore, the partial protection from WT CHIKV infection in AB6 mice suggests other immune factors, such as CD8^+^ T cells also contribute to control of WT CHIKV infection.

EILV/CHIKV-vaccination induced potent and durable CD8^+^ T cell responses against CHIKV structural proteins (Capsid, E3, E2, and E1) at 1 year PV. In comparison, CHIKV 181/25-vaccinated macaques had lower levels of CD8^+^ T cell response reactive to capsid, E3 and E1 at 1 month and only detectable levels of E1-specific CD8^+^ T cell responses at 1 year PV. CD8^+^ T cell epitope specificity can influence infection outcome. For example, CHIKV-specific CD8^+^ T cells preferentially target E1 and E2 proteins upon infection(28, 29). More evidence suggests that CHIKV capsid protein also contains immunodominant epitopes for both B and T cells. Indeed, a low frequency of CHIKV capsid-specific T cells in the aged animals were reported to be associated with virus persistence in CHIKV-LR strain-infected rhesus macaques (30, 31). Vaccines expressing a combination of structural proteins or the entire structural protein of CHIKV have been shown to induce more potent CD8^+^ T cell-mediated responses (28, 32) and this has been used as a strategies to optimize vaccine immunogenicity and increase the potency of cellular and subsequent humoral immune responses (33–35). Thus, a potent, durable and broader repertoires of CD8^+^ T cells contribute to EILV/CHIKV-induced long-lasting protection against CHIKF.

EILV/CHIKV also protected macaques rapidly from CHIKF 6 days post a single vaccination. Transcriptome analysis of PBMCs from EILV/CHIKV-vaccinated macaques at 1 week suggest induction of multiple innate immune pathways, including TLR signaling, type I IFN and IL-12 signaling, antigen presenting cell (APC) activation, NK receptor signaling, Th1, Th2 and B cell signaling pathway. During CHIKV infection, viral RNA is identified by pattern recognition receptors (PRRs), including TLRs 3, 7, and 8, RIG-I-like receptors (RLRs), and melanoma differentiation-associated gene 5 (MDA5). which triggers the synthesis of pro-inflammatory cytokines and type I IFNs (36, 37). Type I IFNs contribute to viral clearance and the control of virus-mediated inflammation, protecting humans and mice against CHIKV infection, and greater expression of IL-12 and IFN-α is correlated with lower viral loads in acute CHIKF patients(38, 39). Cytokine analysis of sera of vaccinated macaques upon WT CHIKV infection suggest increased production of type I IFN and IL-12 in both inactivated WT CHIKV and EILV/CHIKV groups and the latter group had a quicker kinetics of induction. Enhanced levels of inflammatory mediators such as IL-1β, GM-CSF, IL-23 are often associated with CHIKV-induced inflammation responses (38, 40). Here, we noticed induction of these cytokines at an early stage of CHIKV infection in both vaccinated groups, with the EILV/CHIKV group exhibiting faster kinetics. The induction of these inflammatory cytokines in addition to IFN-I and IL-12 are more likely to be associated with NK cell expansion and activation in EILV/CHIKV-vaccinated macaques. Indeed, we observed increased activation of NK cells in both inactivated WT CHIKV and EILV/CHIKV-vaccinated groups and a higher magnitude in the latter group at day 4 PC. NK cells contribute to control of acute CHIKV infection by subset-specific expansion. and dysregulation of NK receptor expression correlates with CHIKV-induced chronic diseases (41, 42). Furthermore, induction of the above innate cytokines could lead to the activation of adaptive immunity. For example, IL-1β and IL-23 act either as signal 3 cytokines for T cell priming or regulating APC function and CD8^+^ T cell responses (43, 44). Consequently, EILV/CHIKV-vaccinated macaques showed higher levels of neutralization titers and strong Th-1 prone immune responses at day 4 PC. Lastly, neutralization antibodies could trigger the antibody-dependent cytotoxicity responses mediated by NK cells and CD8^+^ T cells to control CHIKV infection (45). In summary, EILV/CHIKV-induced type I IFN, NK cell activation, Th-1 responses and neutralization antibodies responses contribute to its rapid protection against CHIKF.

Infection in cynomolgus macaques induces viremia, acute disease signs and viral persistence in some tissues, which are similar to clinical symptoms reported in humans. The cynomolgus macaque has been considered an ideal preclinical model to evaluate the efficacy and safety of CHIKV vaccines (46, 47). In both long- and short-term protection studies, we observed minimal to mild pathological changes in the lymphoid organs of the vaccinated and mock macaques following WT CHIKV infection and the differences between the two groups were minimal. In particular, long-lasting CHIKV disease signs were not observed in the mock group upon WT CHIKV infection as previously reported (48). This is due to the inoculation dose we used in this study, which is directly associated with virological and clinical outcomes (48).

Vaccine produced in mosquito cell line has a risk of causing hypersensitivity reactions due to its potential contamination with mosquito antigens. We previously reported a two-step protocol for improving the purity of EILV/CHIKV. The purified virus was shown to cause minimal histopathology changes and Th-1 cytokine induction in a mouse model of hypersensitivity model (21). Due to its similarities with humans in immune system, anatomy and physiology, guinea pig has been considered to be more similar to humans than between mouse models and humans and thus a more feasible model to assess hypersensitivity(49, 50). Here, we found that EILV/CHIKV did not cause skin hypersensitivity reactions in a guinea pig sensitization model, indicating its favorable safety profile.

Formalin-inactivated CHIKV vaccine is safe and induces robust immunity and strong protection. However, its development was terminated due to high manufacturing costs (51). The live-attenuated 181/25 CHIKV vaccine developed in the 1980s is also highly immunogenic and induces strong protection. However, Phase II clinical trials reported 8% of vaccinees developed arthralgia due to an unstable attenuation (11, 51), which is based on two mutations derived from the WT parent strain with a high risk of reversion that are hard to optimize by genetic engineering (52, 53). Genetic stability is a valid concern for any live attenuated vaccines for CHIKV, including the licensed Ixchiq vaccine.

In summary, the results from this study suggest that EILV/CHIKV is a safe, highly efficacious vaccine, and provides rapid and long-lasting protection in cynomolgus macaques after a single dose via induction of antiviral innate immunity and durable T cell immunity.

## Materials and Methods

### Sex as a biological variable

Both male and female cynomolgus macaques and mice were used in this study. Due to animal availability during the pandemic, single sex (either female or male) macaques were used on the long-term and short-term protection studies respectively. Female guinea pigs were used for skin hypersensitivity study. It’s unknown whether the findings are relevant to male guinea pigs.

### Animal studies

2.5 to 3.5-year-old female and male Mauritian cynomolgus macaques (*Macaca fascicularis*) were purchased from Worldwide Primates, Inc. (Miami, FL). All animals screened negative for TB, SIV, simian T lymphotropic virus, herpes B virus, *T. Cruzi*, SRV, and CHIKV. Six- to seven-week-old C57BL/6 (B6) mice deficient in the IFN-α/β receptor (AB6) were bred and maintained at the University of Texas Medical Branch (UTMB). Cynomolgus macaques were vaccinated intramuscularly (i.m) with_a low dose [LD, 1.3 × 10^6^ plaque forming unit (PFU)] or_high dose (HD, 1.3 × 10^8^ PFU) of the EILV/CHIKV, or 3.1 × 10^5^ PFU of CHIKV 181/25, 5 × 10^5^ PFU of the inactivated wild-type (WT) CHIKV Martinique strain, or PBS. Two weeks before challenge, subjects were bled via the femoral vein. Prior to challenge, each NHP was implanted intraabdominally with a DST micro-T temperature logger (Star-Oddi, Garðabær, Iceland) programed to take a temperature reading every 15 minutes. Macaques were challenged with 1.0 ×_10^5^ PFU of the WT-CHIKV La Réunion (LR) strain at days 6 or 370 post vaccination (PV). For passive immunization studies, AB6 mice were transferred intraperitoneally (i.p.) with diluted sera from mock or vaccinated macaques 24 h before challenge with 1×10^3^ PFU WT CHIKV Martinique strain. Infected animals were monitored daily for clinical signs.

### Virus stocks

EILV/CHIKV was propagated in C7/10 mosquito cells and viral stocks were purified using a two-step procedure as described previously (21). CHIKV LR and Martinique strain stocks were kindly provided by Dr. Trevor Brasel at UTMB. Formalin-inactivation of WT-CHIKV was performed as described previously (20). Briefly, a CHIKV Martinique stock was diluted 1:10 in DMEM media and inactivated by the addition of 20 µl of 10% formaldehyde (Sigma, St. Louis, MO) to a final concentration of 0.1% (v/v), or mock-inactivated by the addition of an equivalent volume of PBS. Virus suspensions were incubated at 37°C with mixing twice daily for 3 days, followed by incubation at 4°C with mixing twice daily for 4 days. Formaldehyde was neutralized by the addition of 3.5% (w/v) sodium metabisulfite (Sigma) to a final concentration of 0.035% (w/v). Inactivation was confirmed by plaque assay.

### Blood collection and peripheral blood mononuclear cell (PBMC) isolation

Blood was collected by femoral venipuncture into EDTA in serum separator/lithium heparin vacutainer tubes (BD Biosciences). Blood tubes were centrifuged at 3000 rpm at 4°C for 15 min first and the upper layer was collected. For PBMC isolation, heparin-treated blood was diluted 1:1 with sterile PBS and carefully layered onto Ficoll-Paque (GE Healthcare Bio-Sciences, Sweden). The tubes were centrifuged at 400 x g for 30 min at 20°C, and then the mononuclear cells were transferred to a sterile centrifuge tube. Cells were washed twice with PBS and resuspend in RPMI medium supplemented with 10% FBS, and 1% l-glutamine (Thermo Fisher). Cells were cryopreserved and stored in freezing medium (10% DMSO/ 90% FBS) in liquid nitrogen tank.

### Plaque assay

Vero cells were seeded in 12-well plates and incubated at 37_°C, 5% CO2 for 16 h. Serum samples were serially diluted in DMEM with 2% FBS and 100 μl of diluted samples were used to infect the monolayers. Plates were incubated at 37°C with 5% CO2 for 1 h. After the incubation, 0.4% agarose overlay medium was added to the infected cells. The plates were incubated at 37_°C with 5% CO2 for 48 h. Plates were fixed with 10% formaldehyde for 30 min. Cells were stained with 0.25% crystal violet for 3 min, then rinsed with H_2_O. The plaques were counted and calculated as PFU /mL.

### Quantitative PCR (Q-PCR)

Blood cells were re-suspended in Trizol (Invitrogen) for RNA extraction. Complementary (c) DNA was synthesized by using a qScript cDNA synthesis kit (Bio-Rad, Hercules, CA). The sequence of the primer sets for CHIKV capsid protein, cytokines and PCR reaction conditions were described previously (21, 54, 55). The PCR assay was performed in the CFX96 real-time PCR system (Bio-Rad). Gene expression was calculated using the formula 2^ ^-[C^t^(target^ ^gene)-C^t^(β*-actin*)]^ as described before (56).

### NanoString sample preparation

Targeted transcriptomics was performed on PBMC samples from macaques. Nonhuman primate (NHP) V2_Immunology reporter and capture probe sets (NanoString Technologies) were hybridized with ∼100 ng of each RNA sample for ∼24 h at 65°C. The RNA:probeset complexes were subsequently loaded onto an nCounter microfluidics cartridge and assayed using a NanoString nCounter SPRINT Profiler. Samples with an image binding density < 0.2 were re-analyzed with a higher concentration of RNA to meet quality control criteria.

### Transcriptional analysis

Analysis was performed on samples at 7 days post-vaccination or 4 days post infection. nCounter .RCC files were imported into NanoString nSolver 4.0 software. To compensate for varying RNA inputs and reaction efficiency, an array of housekeeping genes and spiked-in positive and negative controls were used to normalize the raw read counts. As both sample input and reaction efficiency were expected to affect all probes uniformly, normalization for run-to-run and sample-to-sample variability was performed by dividing counts within a lane by the geometric mean of the reference/normalizer probes from the same lane (i.e., all probes/count levels within a lane are adjusted by the same factor). The ideal normalization genes were automatically determined by selecting those that minimize the pairwise variation statistic and are selected using the widely used geNorm algorithm. The data were analyzed with NanoString nSolver Advanced Analysis 2.0 package for differential expression and to generate principal component analysis (PCA), heatmap, and cell-type trend plots. Human annotations were added for each mRNA to perform immune cell profiling within nSolver and generate the cell-type scores. Normalized data (fold change values and Benjamini–Hochberg adjusted p-values) were exported as an .xlsx file (**Table S1**) and imported into GraphPad Prism version 10.0.1 to generate heatmaps. Figures were prepared using Adobe Illustrator 2024. Any differentially expressed transcripts with a Benjamini-Hochberg false discovery rate (FDR) corrected p-value less than 0.05 were deemed significant unless otherwise stated. Enrichment of differentially expressed transcripts was performed using Ingenuity Pathway Analysis (Qiagen). Samples were filtered for immunity-specific canonical signaling pathways.

### Peptide pools

Fifteen amino acid (aa) peptides overlapping 12 aa spanning the structural proteins of CHIKV were purchased from Sigma and were divided into 4 overlapping peptide pools, including core (peptides 1-53), E3 (peptides 54-77), E2 (peptides 78-211), and E1 (peptides 212-343).

### Intracellular cytokine staining (ICS) and flow cytometry

Cryopreserved PBMCs were thawed and washed once with RPMI medium containing 10% FBS. Cells were incubated with individual CHIKV peptide pools (2μg/ml) for 6 h in the presence of BD GolgiPlug (BD Bioscience). Cells were harvested and stained with antibodies for CD3, CD4, or CD8, fixed in 2% paraformaldehyde (PFA), and permeabilized with 0.5% saponin before adding anti-IFN-γ (e-Biosciences). PBMCs were also stained for the following cell surface markers: CD56 (clone: SP34-2; BD Biosciences) and CD3 (clone: TULY56; eBioscience) for 30 min at 4°C, washed and fixed in 2% PFA. Samples were processed with a C6 Flow Cytometer instrument. Dead cells were excluded on the basis of forward and side light scatter. Data were analyzed with a CFlow Plus Flow Cytometer (BD Biosciences).

### CHIKV IgG ELISA

ELISA plates were coated with 50 ng/well recombinant CHIKV E2 protein (SinoBiological, USA) overnight at 4°C. The plates were washed twice with PBS, containing 0.05% Tween-20 (PBS-T) and then blocked with 2% PBS-T for 1.5 h at 1.5 h at room temperature (RT). Sera were diluted 1:200 in blocking buffer and 50 µl were added per well for 1 h at RT. Plates were washed five times with PBS-T. Goat anti-monkey IgG (Fitzgerald, USA) coupled to horseradish peroxidase (HRP) was added as the secondary antibody at a 1:25000 dilution for 1 h at RT. Color was developed with TMB substrate (Thermo Scientific) and the intensity was read at an absorbance of 450 nm.

### Plaque reduction neutralization assay

Fifty percent plaque-reduction neutralization tests (PRNT50) were performed on Vero cells. Briefly, NHP sera were heat-inactivated at 56°C for 30 min then serially diluted (2-fold dilutions) in 2% FBS MEM media. CHIKV 181/25 stock was diluted to 800 PFU/ml. Then 150 µL of the diluted sera were mixed with 150 µL of virus stock, incubated at 37°C for 60 min. Afterwards, 250 μl of each sera/virus mixture was added to Vero cells, and incubated for 1 h at 37°C, with rocking every 15 min. Then MEM/0.4% agarose overlay media was added to each well, and incubated at 37°C with 5% CO_2_ until plaques appeared (about 48 h). Plates were fixed with 10% formaldehyde and stained with 0.25% crystal violet. The plaques were counted and the PRNT titers calculated as the highest dilution of serum that inhibited 50% of plaques.

### B cell ELISPOT assay

ELISpot assays were performed as previously described (57) with some modifications. Briefly, PBMCs were stimulated with 1µg/ml R848 and 10 ng/ml recombinant human IL-2 (Mabtech In, OH). Millipore ELISPOT plates (Millipore Ltd, Darmstadt, Germany) were coated with CHIKV peptide pools (15 µg/ml), or CHIKV E2 recombinant protein (SinoBiological, USA, 15 ug/ml), EILV/CHIKV (1 × 10^8^ PFU/ well), CHIKV VLP (The Native Antigen Company, Oxford, UK, 15 µg/ml), or anti-human Ig capture Ab (Mabtech). The stimulated PBMCs were harvested, washed, and added in duplicates to assess total IgG ASCs or CHIKV-specific B cells. Plates were incubated overnight at 37°C, followed by incubation with biotin conjugated anti-human IgG (Mabtech) for 2 h at room temperature, then 100 µL/well streptavidin-ALP was added for 1 h. Plates were developed with BCIP/NBT-Plus substrate until distinct spots emerge. After washing, the plates were scanned using an ImmunoSpot 4.0 analyzer and the spots were counted with ImmunoSpot software (Cellular Technology Ltd, Cleveland, OH).

### IFN-γ ELISPOT

Monkey anti-IFN-γ capture Ab pre-coated plates (Mabtech) were washed twice and blocked with RPMI media containing 10% FBS for 30 min. PBMCs were stimulated in duplicates with CHIKV peptide pools (2μg/ml) for 24 h at 37°C. PBMCs stimulated with anti-CD3 (1μg/ml, e-Biosciences) or medium alone were used as controls. This was followed by incubation with biotin-conjugated anti-IFN-γ for 2 h at room temperature, and then conjugated streptavidin-HRP for 60 minutes. Plates were developed with TMB substrate, followed by washing and scanning using an ImmunoSpot 4.0 analyzer. The spots were counted with ImmunoSpot software (Cellular Technology Ltd, Cleveland, OH) to determine the spot-forming cells (SFC) per 10^6^ PBMCs.

### Cytokine bead arrays

Sera of infected NHPs were inactivated with 5 megarads Mrads in a Cobalt-60 irradiator at Galveston National Lab (GNL, UTMB) before used for cytokines, chemokines, and growth factors measurement using LegendPlex bead-based multiplex technology and NHP inflammation 13-plex kits (BioLegend). Serum samples were processed in duplicate at a 1:4 dilution according to the manufacturer’s instructions. Following bead staining and washing, at least 3,900 bead events were collected on an Aurora cytometer (Cytek) using SpectroFlo software. The raw .fcs files were exported and analyzed with BioLegend’s cloud-based Qognit data analysis software. The concentration data based on standard curves were exported to Microsoft Excel to calculate average fold-change values over the mock group average baseline.

### Statistical analysis

Survival curve comparison was performed using GraphPad Prism software 10.0.1, which uses the log-rank test. Values for viral load, cytokine production, antibody titers, memory B cell frequency, and T cell responses were compared using Prism software (GraphPad) statistical analysis and were presented as means ± SEM. *P* values of these experiments were calculated with a non-paired Student’s t test.

### Study approval

All animal studies were approved by the Institutional Animal Care and Use Committee at UTMB (protocol # 1704031A and #1904041).

## Data and materials availability

All data are available in the main text or the supplementary materials.

## Acknowledgements

This study was supported by National Institute of Health grants R01AI127744 (T.W.), R01 NS125778 (T.W.), and R01 AI176670 (T.W.). H.L. was a recipient of a summer internship from NIAID T35 training grant (AI078878, PI: T.W.). We thank Drs. Elena Frolova and Ilya Frolov for providing EILV/CHIKV stocks, Ms. Grace Rafael and Yunmei Yun for technical assistance, Terry Juelich for preparation of inactivated sera samples for cytokine analysis and Dr. Linsey Yeager for assisting in manuscript preparation. We also thank Drs. Doug Brining, John Lowry, Andrew Kocsis, Erin Lee, Thomas Geisbert, and Chad Mire for their constructive suggestions of the study and the technical support of the animal staff team at the Galveston National Laboratory.

## Author contributions

A.A., C.W., H.L., K.P., L.R., D.M., and Y.C. performed the experiments. A.A., C.W., S.W., and T.W. designed the experiments. Y.C., A.W.E.F., S.T., and J.C. provided critical reagents and supplies, A.A., C.W., H.L., S.W., L.R., D.M., J.C., and T.W. analyzed the data. A.A., C.W., and T.W. wrote the initial draft of the manuscript and other coauthors provided editorial comments.

## SUPPLEMENTARY FIGURE LEGENDS

**Supplementary Figure 1. Viremia and histopathology of CHIKV-infected macaques 1 year post vaccination. A-B.** Viremia was measured by Q-PCR at indicated days post-challenge (PC). ** *P* < 0.01 or * *P* < 0.05 compared to mock group. Unpaired, 2-tailed Student’s t test was used to determine the differences. Data are presented as means ± standard error of the mean (SEM). **C.** Representative images of H& E-stained spleens of vaccinated macaques at day 14 PC. Comparatively, minimal to mild decreased lymphocytes in lymphoid nodules (arrows) were observed in some animals.

**Supplementary Figure 2. CHIKV-specific T cell responses in vaccinated macaques. A.** PBMCs of day 30 vaccinated NHPs were cultured *ex vivo* with CHIKV capsid, E3, E2 and E1 peptide pools for 6 h, and stained for IFN-γ, CD3, CD4, or CD8. Total number of IFN-γ^+^ CD4^+^ and CD8^+^ T cells is shown. **B.** ELISPOT quantification of peripheral T cell responses. PBMCs of day 7 and day 30 vaccinated NHPs were stimulated with CHIKV capsid, E3, E2 and E1 peptide pools for 24 h. Spot forming cells (SFC) were measured by IFN-γ ELISPOT. Data are shown as # of SFC per 10^6^ cells. \*\**P* < 0.01, or \**P* < 0.05 compared to mock. ^###^*P* < 0.001, ^##^*P* < 0.01, or ^#^*P* < 0.05 compared to CHIKV 181/25.

**Supplementary Figure 3. CHIKV specific memory B cell and antibody responses in EILV/CHIKV-vaccinated NHPs in long-term protection study. A-B.** CHIKV-specific memory B cell (MBC) responses by ELISPOT analysis. PBMCs of day 30 (**A**) and day 7 (**B**) vaccinated NHPs were stimulated for 7 d with R848 plus rIL-2 and seeded onto ELISPOT plates coated with (**A**) CHIKV VLP and EILV/CHIKV or (**B**) CHIKV capsid, E3, E2 and E1 peptide pools. The frequencies of CHIKV-specific ASCs per 10^6^ input cells in MBC cultures from the subject. **C.** Neutralizing activity against CHIKV 181/25 in sera collected at various time points PV was measured by PRNT. ^##^*P* < 0.01, or ^#^*P* < 0.05 compared to CHIKV 181/25.

**Supplementary Figure 4. Hypersensitivity assay in a guinea pig model.** Guinea pigs were sensitized by exposure to female *Ae. albopictus* mosquitoes four times during a 14-day period. On day 43, animals were injected i.d. on footpad with 20µg SGE protein, 1.3 × 10^8^ PFU of EILV/CHIKV or PBS (mock). 30 min to 2 h after challenge, skin reactions were assessed. **A.** Study design. **B.** Representative image of skin reactions at 2 h post injection on day 43. **C.** Skin swelling was measured via 30 min post inoculation. \*\*\**P* < 0.001 compared to mock. ^##^*P* < 0.01 compared to SGE group.

**Supplementary Figure 5. EILV/CHIKV induced innate cytokine responses.** NHPs were vaccinated with a LD EILV/CHIKV (n =4), CHIKV 181/25 (n =3) or PBS (mock, n= 3). **A-E.** On days 1, and 2 PV, blood cytokines levels were determined by Q-PCR assay. Data are presented as means ± SEM of fold increase compared to the mock group. ^#^*P* < 0.01 compared to CHIKV 181/25.

## Supplementary Methods

### Histopathology

At day 14 PC, spleens were collected from all animals scheduled for euthanasia and placed in 10% formalin (Thermo Fisher Scientific, Waltham, MA, USA) before embedment in an optimal cutting temperature compound. H&E staining was performed at the Histopathology Laboratory Core of UTMB. Slides were examined by a board-certified pathologist at the Experimental Pathology Laboratories, Inc. (EPL^®^) in Sterling, Virginia.

### *Aedes albopictus* salivary gland protein extract

Around a hundred salivary glands from 6-day old sugar-fed *Ae. albopictus* females, (strain: Lake Charles, LA) were dissected in droplets of PBS as described before (1). Dissected salivary glands were then combined and ground in protein lysis buffer (50 mM Tris-HCl, pH 8.0, 150 mM NaCl, 1 mM EDTA, 1% Triton X-100). The resulting salivary gland extract was kept on ice and for storage frozen at – 80 C.

### Guinea pig sensitization study

6 to 8-week-old female Dunkin Hartley albino Guinea pigs were purchased from Charles River (Wilmington, MA). Animals were sensitized with 50 females *Ae. albopictus* mosquitoes for 4 times on a 14-day interval. On day 43, guinea pigs were injected intradermally (i.d.) at the mosquito probing/feeding area with 20 µg *Ae. albopictus* SGE, 1.3x 10^8^ PFU of purified EILV/CHIKV, or PBS-G (mock). 30 min to 2 h following injection, inoculation sites were assessed via measurements of site swelling and imaging.

