## Supplementary figures for "A safe insect-based Chikungunya fever vaccine affords rapid and durable protection in cynomolgus macaques"

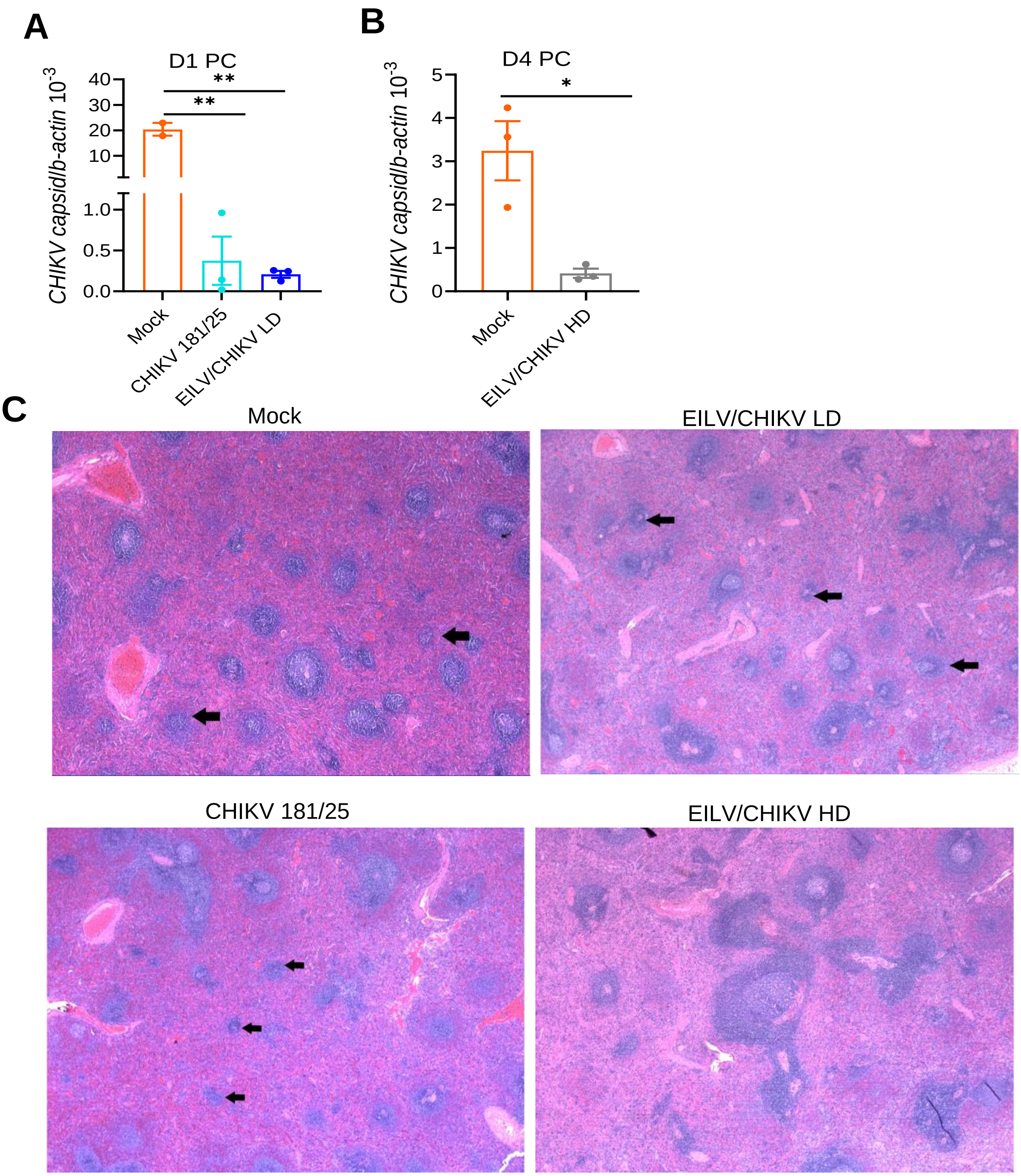


**Supplementary Figure 1**

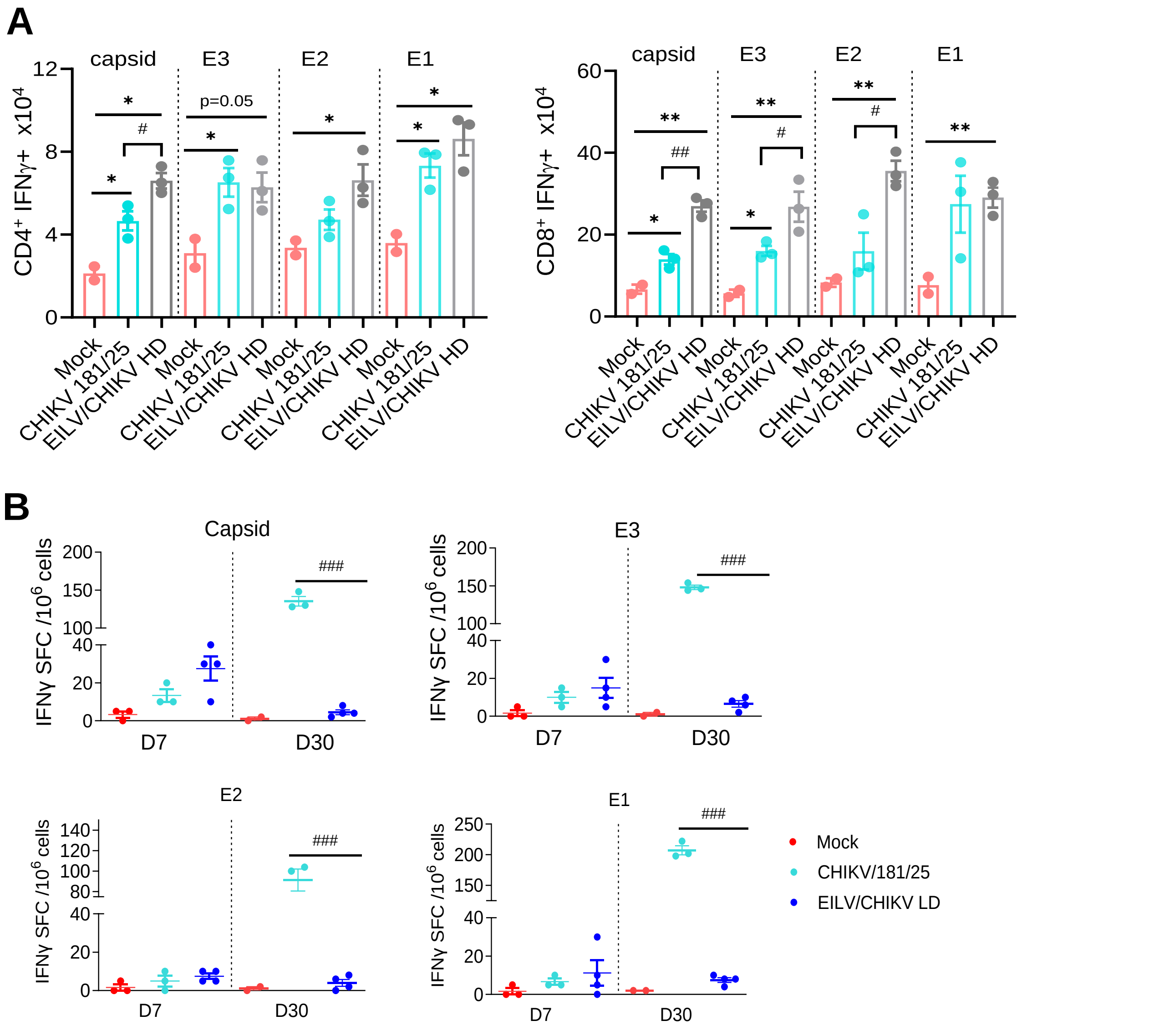


**Supplementary Figure 2**

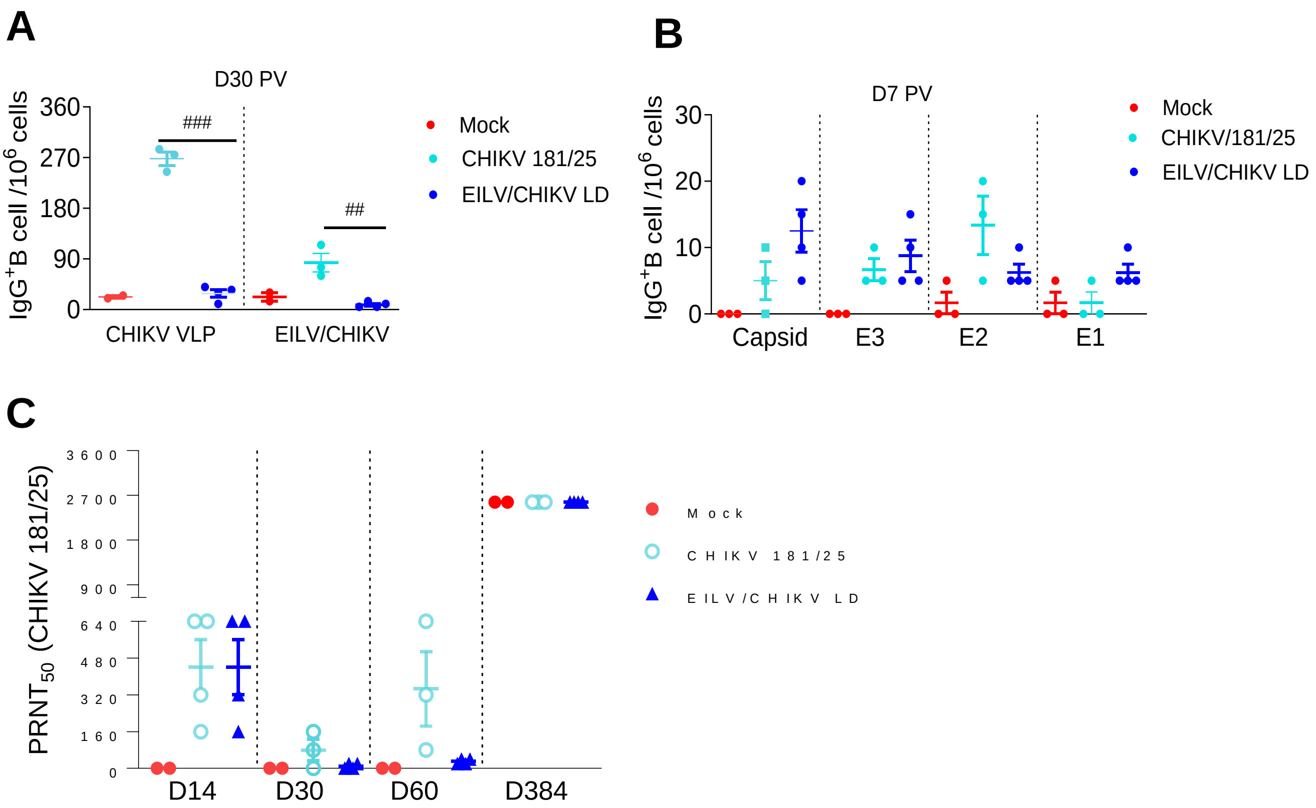


**Supplementary Figure 3**

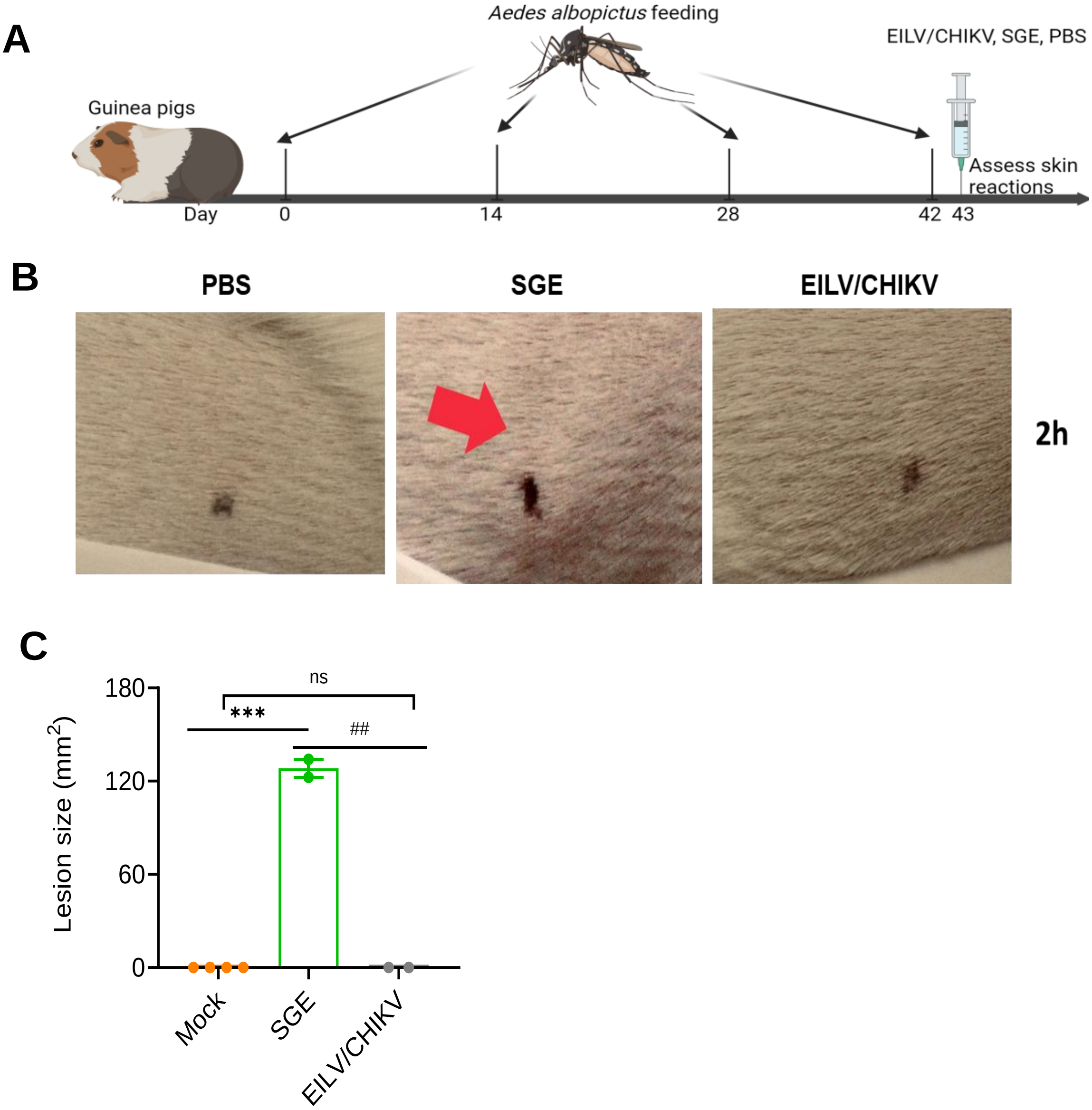


**Supplementary Figure 4**

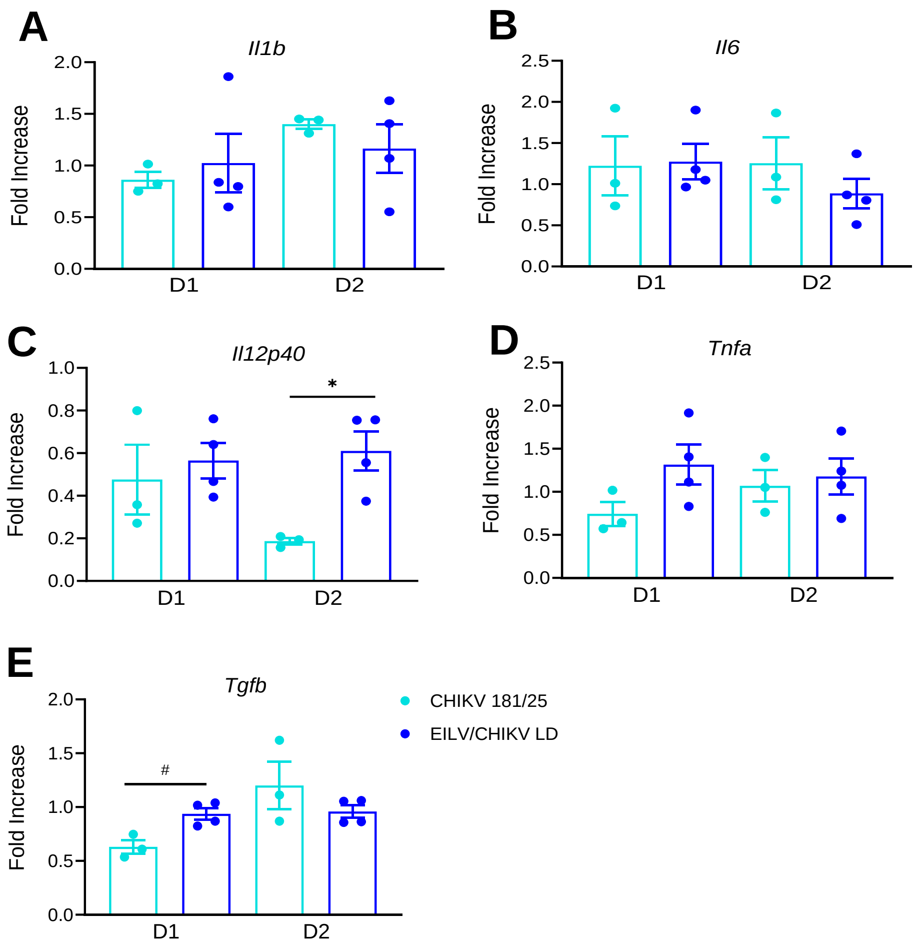


**Supplementary Figure 5**
